# Evolutionary history and origin of Tre1 superfamily shed light on its role in regulating blood brain barrier

**DOI:** 10.1101/2021.04.29.441935

**Authors:** Norwin Kubick, Pavel Klimovich, Irmina Bieńkowska, Mariusz Sacharczuk, Michel-Edwar Mickael

## Abstract

Understanding how the evolutionary relationship between immune cells and the blood-brain is important to devise therapeutic strategies that can regulate their critical function. In vertebrates, immune cells follow either a paracellular or transcellular pathway to infiltrate the BBB. In drosophila glial cells form the BBB that regulates the access of immune-like cells to the drosophila brain. However, it is still not known which route immune-like cells follow to infiltrate the drosophila brain. In vertebrates, paracellular migration is dependent on PECAM1, while transcellular migration is dependent on the expression of CAV1. Interestingly drosophila genome lacks both genes. Tre1 superfamily (Tre1, Moody, and Dmel_CG4313) play a diverse role in regulating transepithelial migration in drosophila. However, its evolutionary history and origin are not yet known. We performed phylogenetic analysis, together with HH search, positive selection, and ancestral reconstruction to investigate the Tre1 family Interestingly we found that Tre1 exists in mollusks, insects, ambulacria, and sclaidphora. Moody is shown to be a more ancient protein and it existed since cnidaria emergence and has a homolog (GPCR84) in mammals. The third family member (Dmel_CG4313) only exists in insects. The origin of the family seems to be related to the rhodopsin-like family and in particular family α. We found that opsin is the nearest receptor to have a common ancestor with the Tre1 superfamily that seems to have diverged in sponges. We investigated the positive selection of the Tre1 family using PAML. Tre1 seems to have evolved under negative selection, whereas Moody has evolved during positive selection. The sites that we found under positive selection are Likely to play a role in the speciation of function in the case of Moody. We have identified an SH3, in Tre1 and, moody and Dmel_CG4313. Sh3 is known to play a fundamental role in regulating actin movement in a Rho-dependent manner. We suggest that Tre1 could be playing an important role in paracellular diapedesis in drosophila.

## Trans-epithelial migration

The blood brain barrier (BBB) functions to insulate the central nervous system from its changing molecular environment ^1^. This insulation is essential to defend neurons from toxic substances and to maintain invariable ionic composition within the nervous system to enable efficient neuronal conductivity. In the vertebrate central nervous system, BBB function depends on an efficient separation between the blood components and the nervous system and is maintained by the sealing of brain endothelial cells by specialized tight junctions. However, insulation of a vertebrate’s peripheral nervous system is performed by myelinated Schwann cells that produce a myelin sheath covering the axon in segments separated by the nodes of Ranvier. The segmented myelin sheaths enable saltatory movement of the nerve impulse from node to node. The Schwann cell in each myelinated segment forms a continuous array of septate junctions with the underlying axon. These junctions insulate the Na+ channel–rich domain from the axon segment that undergoes myelination ^2^. The *Drosophila melanogaster* BBB is produced by the sealing of the nervous system from the outside surrounding environment by a layer of large glia cells called subperineurial glia (SPG); these cells tightly adhere to each other using an array of septate junctions formed at the lateral borders of these cells. Similar to vertebrates, sealing of the *Drosophila* nervous system is essential to its protection from toxic substances and high potassium ion concentrations characteristic of the surrounding hemolymph. In addition to the membrane proteins NeurexinIV, Neuroglian, and Contactin, *Drosophila* septate junctions contain components of vertebrate tight junctions such as the claudin-like proteins. In the fly, septate junctions functionally replace vertebrate tight junctions to provide an epithelial barrier in various tissues, including the ectoderm, salivary gland, tracheal cells, and glia cells ^3^.

Both Tre1 and Moody are expressed in the drosophila blood brain barrier. Tre1 superfamily includes three homologs; Moody, Tre1 and CG313 [ref]. It has been shown that in moody mutant flies, the BBB was compromised and drosophila manifested behavior abnormalities. Interestingly, large spaces between the SPG cell junctions, furthermore, *moody* mutant SPG cells exhibit abnormalities in the actin cytoskeleton [ref]. However, the molecular link between Moody signaling, actin skeleton arrangement, and the establishment of elongated septate junctions along the SPG plasma membrane is not clear. Tre1 has been shown to play an important role in transepithelial migration. Tre1 was proposed to be a trehalose receptor^4 5 6^. However it was found that the proposed function of Tre1 is done by a neighboring gene (e.g., Gr5a)^5^. It was found that Tre1, that is essential for Drosophila germ Transepithelial migration ^7^. Interestingly it was shown that inhibiting small Rho GTPases in germ cells affects transepithelial migration, suggesting that Tre1 signals through Rho1. It was proposed that Tre1 acts in a manner similar to chemokine receptors required during transepithelial migration of leukocytes, implying an evolutionarily conserved mechanism of transepithelial migration. Recently Tre1 has been identified in the drosophila BBB [ref]. However its role in is still mysterious.

The evolutionary history and origin of Tre1 is not yet known. Tre1 has been reported in drosophila^7^. However if it exists outside Arthropoda family is not yet known. It is closely related to 7 transmembrane GPCRs^7^. Namely it was shown that there is high similarity between Tre1 and melatonin as well as histamine, and serotonin receptors. It was also suggested to be related to chicken melatonin receptor, mouse neuropeptide Y receptor, human dopamine receptor, and scallop Go-coupled rhodopsin^4^. However, The exact GPCR family is not yet known. Additionally, Tre1 origin and evolutionary history are also not yet known. If it was subjected to a negative selection that allows it unique function in insects has not been investigated. Moody expression has been studied extensively in drosophila. Its structure has been identified to follow GPCRs structure ^8^. However the exact family where moody belong is still not clear. A high similarity has been proposed between GPCR84 and Moody. However if they belong to the same family is still unknown. Furthermore DMEL_GC313 evolutionary expression is also not known. Additionally Tre1 family in transmigration of hemtocytes into the drosophila is still not known.Taken together these observation highlight for studying the history of three family members, to consider their origin, positive selection and structure.

In this research we employed phylogenetic analysis to assess the origin and the evolutionary history of Tre1 super family. In particular, we preformed multiple sequence alignment, tree phylogenetic building, ancestral sequence construction. We then used Blastp and HHSearch to infer the nearest protein that could be related to the Tre1 super family (Tre1, Moody, Dmel_CG4313). After that we investigated the positive selection structural similarity and motif identification in the three families as well as in the originals sequence. We found that That Moody is the most ancient and most continuously expressed genes of the family, being expressed from cnidarian to humans and being subjected to global positive selection. Tre1 was only expressed and identified in invertebrates. Dmel_CG4313seems to be only in insects. The origin of the family seems to have diverged from a common ancestor of rhodopsin. These results indicate that Tre1 family belongs to the rhodopsin like (α family), as they have they express the NPXXY motif at H7. We identified an SH3 motif that is expressed in Tre1 and Moody. SH3 is known to interact with Rho in paracellular in vertebrates. We postulate that Tre1 and Moody could be performing a regulatory role analogues to PECAM1 vertebrates by adjusting cell adhesion in the glial layer of the drosophila melanogaster, in order to regulate the infiltration of hematocytes.

## Methods

### Database search

The focus of this research was investigating the relationship between Tre1s’ molecular evolution and their functions. Due to the diverse nature and long evolutionary history, we studieded the protein sequences rather than the DNA sequences could be more informative. Moreover, to make sure that our analysis is a reasonable representation of Tre1s’ evolutionary history, we chose 12 species that span more than 500□million years. Drosophila Tre1 protein family was used for BLASTP searches against proteomes of Western gorilla (Gorilla gorilla), Common rat (Rattus norvegicus), House mouse (Mus musculus), Carolina anole (Anolis carolinensis), red junglefowl (Gallus gallus), African clawed frog (Xenopus laevis), Zebrafish (Danio rerio), sea squirt (Ciona intestinalis), common fruit fly (Drosophila melanogaster), sea anemone (Nematostella vectensis), and Hydra (Hydra vulgaris). Sequences were selected as candidate proteins if their E values were ≤1e-10. Sequences were further filtered for having 3 Ig-like domains using CDD^9^.

### Alignment and phylogenetic analysis

Phylogenetic investigation was done in 3 steps. First, Tre1 family amino acid sequences were aligned using MAFFT via the iterative refinement method (FFT-NS-i). After that, we employed ProtTest to determine the best amino acid replacement model. ProtTest results based on the Akaike information criterion (AIC) suggested that the best substitution model is LG+I+G+F, where LG is the substitution model supplemented by a fraction of invariable amino acids (‘+I’) with each site assigned a probability of belonging to given rate categories (‘+G’) and observed amino acid frequencies (‘+F’). The third step included using the protein alignment and the resulting substitution model, in applying 2 different phylogenetic methods to construct the tree, namely, (1) maximum likelihood and (2) Bayesian inference. We performed the maximum likelihood analysis using PHYML implemented in Seaview with 5 random starting trees. We applied Bayesian inference analysis using MrBayes where we implemented a Markov chain Monte Carlo analysis with 1000,000 generations to approximate the posterior probability and a standard deviation of split frequencies <0.01 to indicate convergence as previously described.

### 2.3. Ancestral Sequence Reconstruction (ASR)

We applied the maximum likelihood method to infer the ancestral sequence of each of the proteins investigated. For each protein, we used the ASR algorithm implemented in MEGA6 to build ancestral sequences ^10^. This was followed by BlastP against the nearest earlier diverging organism. BlastP outcome was only accepted if the E□value threshold was less than e-10.

### HHsearch

HHsearch method was used to examine the evolutionary history of Tre1. Only proteins that have already diverged before Tre1 were considered as candidate parents ^11^.

### Positive selection analysis

To determine whether members of the Tre1 family underwent positive selection during evolution, a maximum likelihood approach was employed^12^. In the first instance, respective complementary DNAs (cDNAs) were downloaded for each of the Tre1 protein sequences and aligned according to their codon arrangement. Next, we investigated positive selection using CODEML which is part of PAML v4.4 program suite. We used substitution rate ratio (ω) of nonsynonymous (dN) to synonymous (dS) mutations as an indicator of the selection type. The models used were basic, branch, branch-site, and sites models. The basic model calculates a global ω ratio averaged over all sites and all lineages in the tree. The branch model permits ω ratio to vary among branches in the phylogeny. Using model 2 option, we calculated 2 ω values, the first is specific for the investigated branch in the tree and the other value is for the rest of the branches in the tree. Branch-site models allow ω to vary both among sites in the protein and across branches on the tree. Branch-site models detect positive selection affecting only a few sites along the tested branches. For site models, 2 tests were performed, namely, (M1a versus M2a) and (M7 versus M8). The nearly neutral model (M1a) covers sites under purifying selection (0□<□dN/dS□<□1) as well as sites under neutral evolution (dN/dS□=□1), whereas the positive selection model (M2a) includes sites that evolved under positive selection (dN/dS□>□1). For each model investigated, a likelihood ratio test (LRT) was performed between the respective model and its neutral counterparts. P values were calculated for the χ2 test according to the following equation; P value□=□χ2 (2*Δ(ln(LRTmodel)□-□ln(LRTneutral)), number of degrees of freedom).

### Linear motifs, GPI prediction, metalloproteinase cleavage site prediction

To investigate Tre1s’ molecular evolution-function relationship, we used 3 approaches. First, we searched the protein sequences of the Tre1s for linear motifs. These motifs are composed of short stretches of adjacent amino acids that act as putative protein interaction sites. We performed the search using the ELM server http://elm.eu.org/ with a motif cut-off value of 100^13^. In the second analysis, putative GPI anchoring positions were predicted using GPI prediction tool http://mendel.imp.ac.at/gpi/cgi-bin/gpi_pred.cgi where a search was performed to locate cleavage site residues that could anchor sites for the GPI signal as described in the work by Pierleoni^14^. Third, we used the server https://prosper.erc.monash.edu.au to detect putative metalloproteinase cleavage sites.

### Functional specificity

We employed SDPpred to identify specificity-determining positions (SDPs) in Tre1s’ multiple alignment sequences. The SDPs are amino acid residues that could be responsible for the functional specificity of Tre1 proteins^15^. SDPpred was used to compute a Z score for each alignment column. Z score is an indicator of the probability that the position is a real specificity determinant.

The Bernoulli estimator was used within SDPpred to create a recognition cut-off (B-cut-off) to assess the significance of the Z scores. Positions with a score value larger than B-cut-off were accepted to predict higher specificity probability

### Functional divergence

Type I and II functional divergence between gene clusters of the Tre1 family was estimated through posterior analysis using the DIVERGE v2.0 program^16^. Functional type I divergence determines amino acid that are highly different in their conservation between 2 groups, indicating that these residues have undergone altered functional constraints. The clusters were pairwise compared. The coefficient of type I functional divergence (θI) was calculated between members of all pairs of clusters. θI values significantly larger than 0 indicated site-specific altered selective constraints or shifts in amino acid physiochemical properties that might have resulted by gene duplication and/or speciation. [add type II divergence] [rewrite]

## Results

### Alignment and phylogenetic tree

Phylogenetic investigations identified Tre1 members only in invertebrates. We also found only 1 homolog Ambulacraria (Echinodermata, Patiria_miniata) that was separated from Chordata around 600 MYA as well as various protostomia that emerged around 610 MYA including Scalidophora (animal name), Panarthropoda (animal name). In Mollusca, we identified Tre1 homolog in Bivalves (Mizuhopecten yessoensis, PECTIN_MAXIMUS, Crassostrea gigas, Crassostrea virginica) but not in Monoplacophorans. We found one homolog in Cephalopods (octopus sinensis) but not in Scaphopods or Aplacophorans or Polyplacophorans (chitons) or Annelida. Tre1 also does not seem to have any homologs in Placozoa or Cnidaria, Ctenophora, porifera or choanoflagellates. No homologs for Tre1 was found outside kingdom Animalia. (figure 1c). For CG4322 (moody), we were able to find it in Placzoa, cniadria, Ambulacraria, Scalidophora, panathropdoa and mollusca (expand two/three sentences).

**Figure 1.**
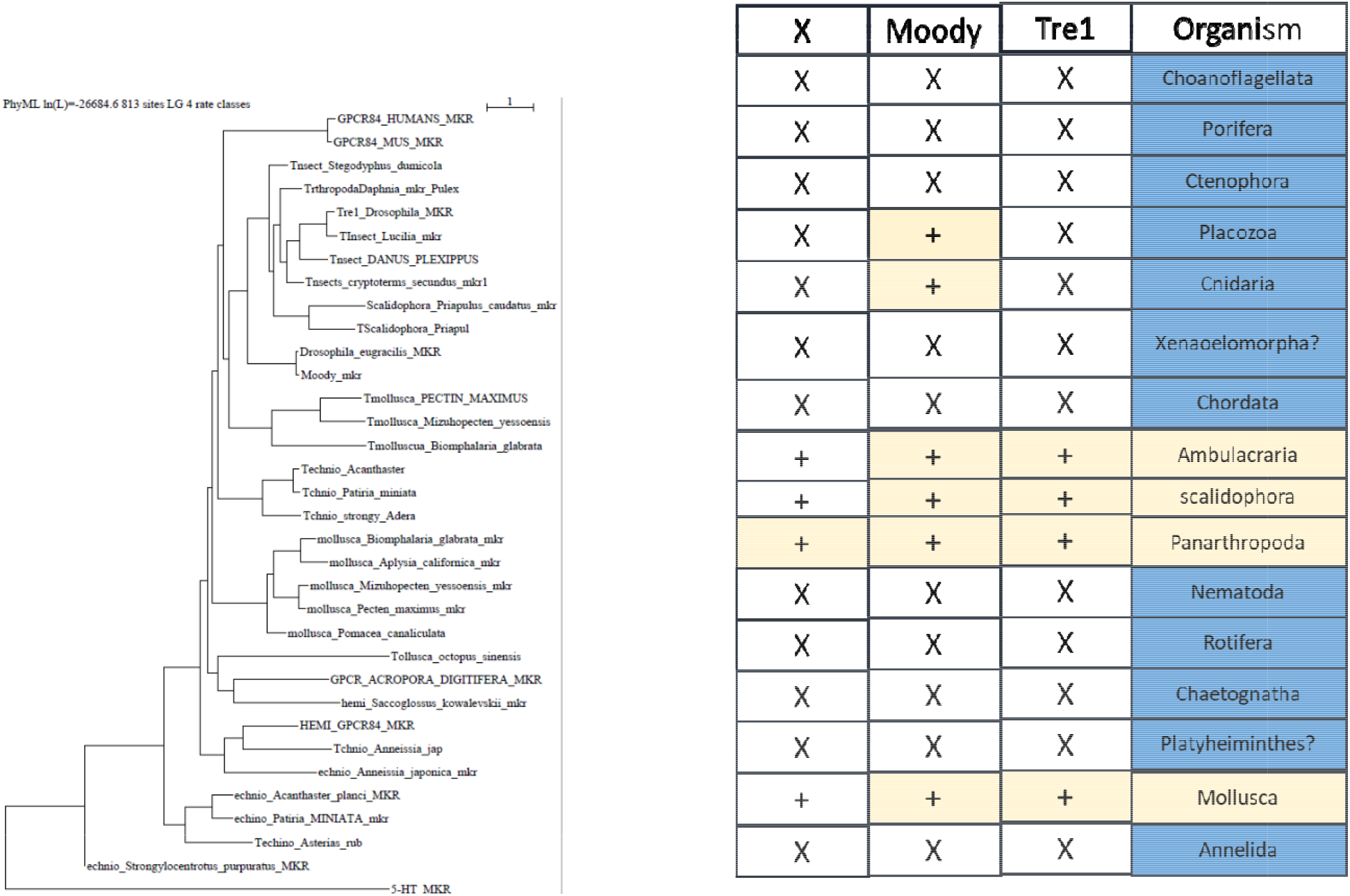
Evolutionary histroy of the Tre1 family. High similarity between Tre1 orthologs (%) (check tree)

### Structural analysis

The Tre1 sequences had a high degree of similarity on different levels, including structural, motif, GPI function, as well as specificity of their residues. On the structural level, all Tre1 family members shared 3 Ig-like domains of the C2 type, as highlighted in the alignment. We also identified several motifs that were shared by more than one Tre1 group. Motif KVTVNYP that acts as SH3 recognition domain was detected in NTM, OPCML, and LSAMP. Motif SEVSNGT/SNGTSRR, which serves as putative CK1 phosphorylation site, was located in LSAMP and invertebrates. In addition, motif TNASIM/SNGTSR that acts as N-glycosylation was detected in Tre15 and NTM. With respect to GPI anchor positions, we were able to locate putative GPI anchoring sites in all Tre1 family members. OPCML and NTM shared several putative metalloproteinase cleavage sites such as V-50, H-94, as well as P-187. This is also supported by the observation of putative GPI-linked anchors in all Tre1 members.

### Functional specificity

We employed SDPpred to determine positions that were well conserved within Tre1 groups but differ between them. The defined amino acid residues might be causing differences in protein functional specificity as well as correct recognition of interaction partners. We observed that these residues could be grouped into 2 locations: location 182 (second Ig-like C2) and 226 (third Ig-like C2). For location 182, LSAMP had (E), OPCML, NTM, and NEGR1 had (D), whereas Tre15 and invertebrates had (G). For residue number 226, LSAMP, OPCML, NTM, and Ciona intestinalis had (s), whereas Tre1 5 had (M) and NEGR1 (A).

### Positive selection investigation

Our results suggested that Tre1 has a large variation with respect to evolutionary selection. We employed global, branch, branch-site, and site models in the CODEML program of PAML v4.4 to examine whether members of the Tre1 family underwent positive selection. For the global model, our results indicate that Moody was under positive selection with ω value of 1.18 and P□<□.001. There was little detected variation among branches with Arthropoda (moody) under insignificant positive selection, while humans(GPCD84, Arthropoda (Tre1) and Arthropoda (Moody)) where under in significant positive selection [are we going to add branch site?]

## Discussion

Analysis of the Tre1 family from a phylogenetic perspective provided the basis for understanding its functional diversity. Phylogenetic analysis was conducted to trace the evolutionary history of the Tre1 family in 12 species. Positive selection, functional specificity, and functional divergence were analyzed at the amino acid level to investigate the evolutionary drivers of the Tre1 family function. We also investigated GPI putative sites, metalloproteinase sites, and motifs. We found three homologs of Tre1, namely, Tre1, Moody, and CG313. Moody first diverged during cnidaria emergence and it is expressed in mammals. Tre1 has a shorter evolution history as it first appeared in Nephrozoa in both Ambulacraria and Protostomia (Ecdysozoa and Spiralia) but not in Chordata. CG313 is only found in insects. We found that the nearest genes that could constitute a putative origin for the Tre1 family are opsin, olfactory receptors, and GPCR161. However, Blastp and structural analysis show that the nearest candidate to have a common ancestor with the Tre1 family is opsin. On an individual family level, we found Tre1 melatonin and Moody are nearest to melanopsin. Positive selection analysis showed that only the Moody family was under positive selection. The rest of the tre1 was shown not to be under positive selection. This suggests that the function of Tre1 and Moody could be tightly conserved among the species that express it. On a structural level, the Tre1 family consists of 7 transmembrane with the main motif of the rhodopsin-like family (NPY). The Tre1 super family shares a trysin sorting signal protein that interacts with the mu subunit of the adaptor signal to transmit the signal to (where?).They also express the SH3 that was show to play an important role in BBB integrity. [other structural difference] The Tre1 is mostly expressed in the BBB and is likely to represent a step toward complexity in controlling access of nutrients and cells to the brain (how that related to the findings).

### Evolution history of the Tre1 family (origin, expression)

Tre1 has a diverse evolutionary history in invertebrates as well vertebrates. Our phylogenetic analysis suggests that Tre1 family (Tre1,Moody and CG313) has first appeared in cnidaria. Investigating the ancestral sequence of Tre1 showed that the nearest sequence for it to be melatonin, somatostatin and 5-HT receptor. The ancestral sequence of Moody includes melonopsin, free fatty acid and red sensitive opsin (figure 2). Tre1 super family does not have any similarity with previously suggested families such as trehalose (ref). Tre1 super family seems have to diverged from a common ancestor of a rhodopsin like transmembrane protein. The ancestral sequence of the Tre1 super family is similar to opsin, olfactory receptor, and GPCR161 (figure 2). Investigating the common motif and similarity (figure 3), suggest that opsin is higher probable candidate to be the nearest protein diverging within the ancestral sequence. This is also supported by Blastp (e value). These findings are in agreement with previous report that linked rhodospin-like family to opsin^17^. It has been suggested that in metazoan, the first opsin originated from the duplication of the common ancestor of the melatonin and opsin genes in a metazoan (Placozoa plus Neuralia) ancestor^18^. Other reports suggested a link between fungi and metazoan opsin^17^ that first appeared 1300 Mya. Whether this ancestral protein was capable of processing light is still not clear. Interestingly, it has been suggested that dipterans possess an ancestral set of five core opsins which have undergone several lineage-specific events including an independent expansion of low wavelength opsins. If any of Tre1 super family could be implicated in light sensitivity is still unknown. Interestingly histamine was reported to activate Tre1 receptor^19^. Whether it is the only GPCR ligand to be able to bind to Tre1 receptor is unknown. We can postulate that various ligands could be able to activate Tre1 due to the promiscuous nature of receptor in lower inevrtabets^20^.

**Figure 2.**
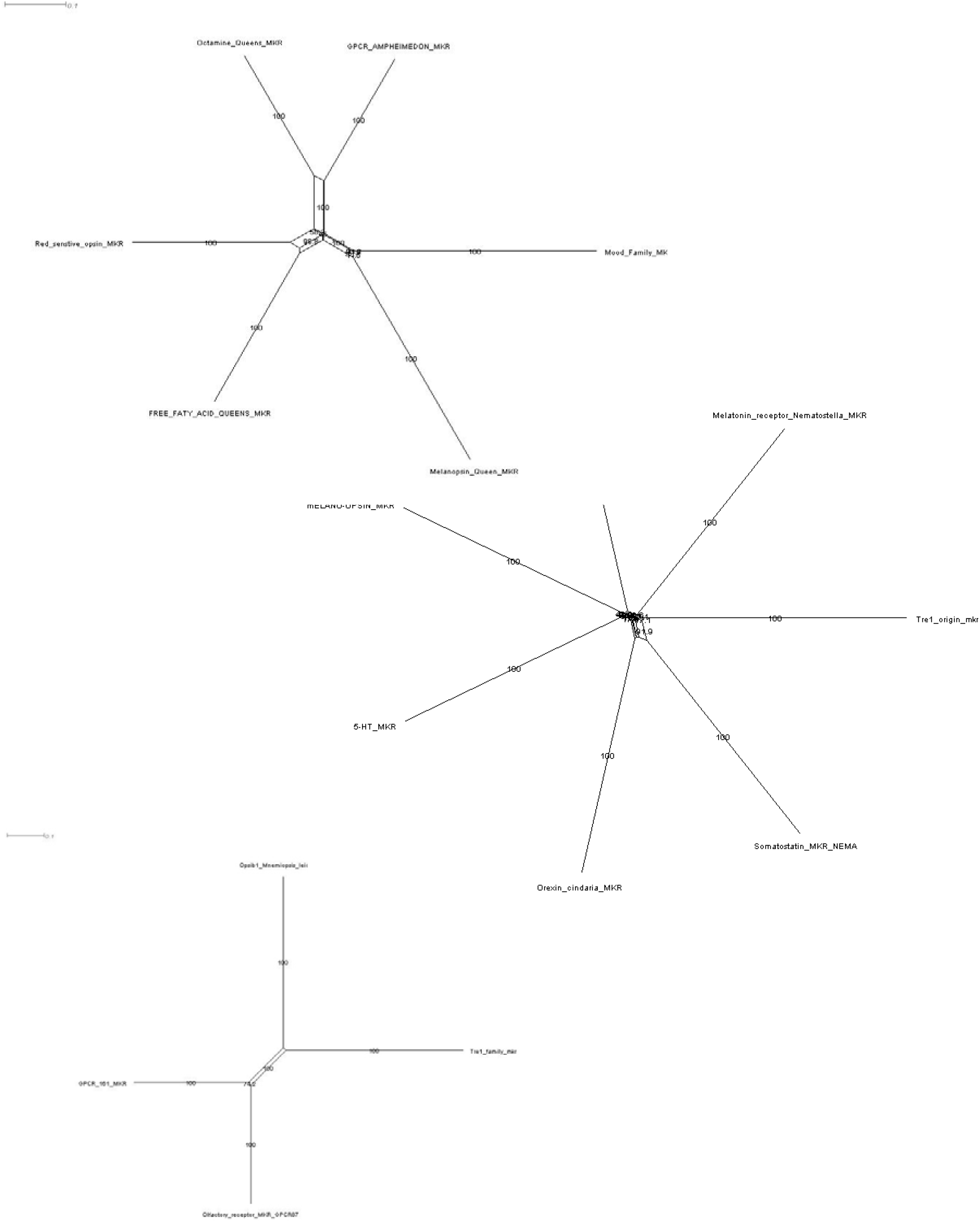
Ancestral sequence construction and network tree for Tr e1. Tre1 could have diverged from GPCR_LIKE Moody or melatonin or EK arthoopsin. (update) (nodes names are unclear).

**Figure 3.**
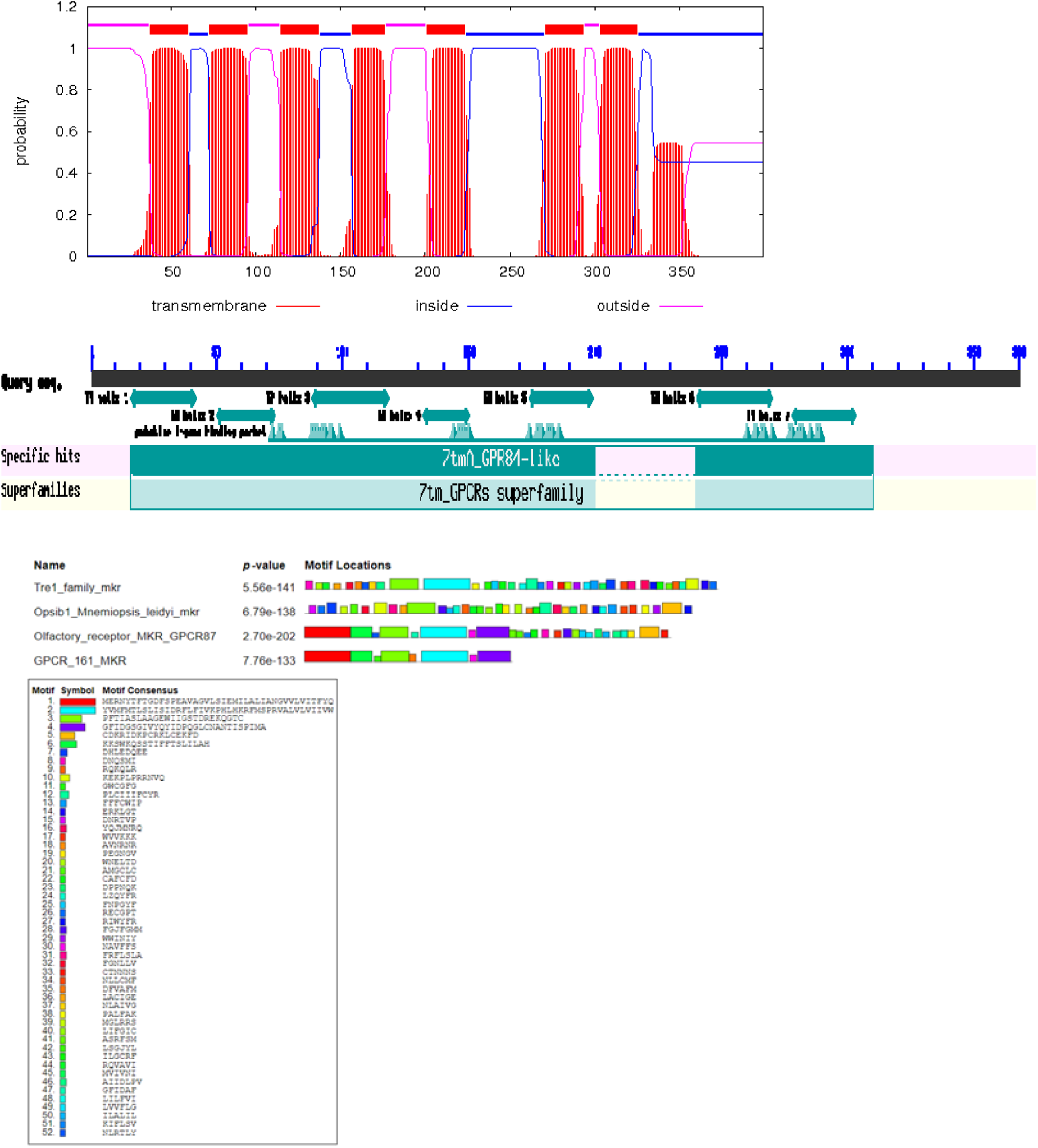
Tr e1 in animals structure. Upper panel shows Tre1 in drosophila with 7 predicted transmembrane regions common in GPCRs. (may be add main here, tre1, and moody and the third separately in supplementary)

**Figure 4.**
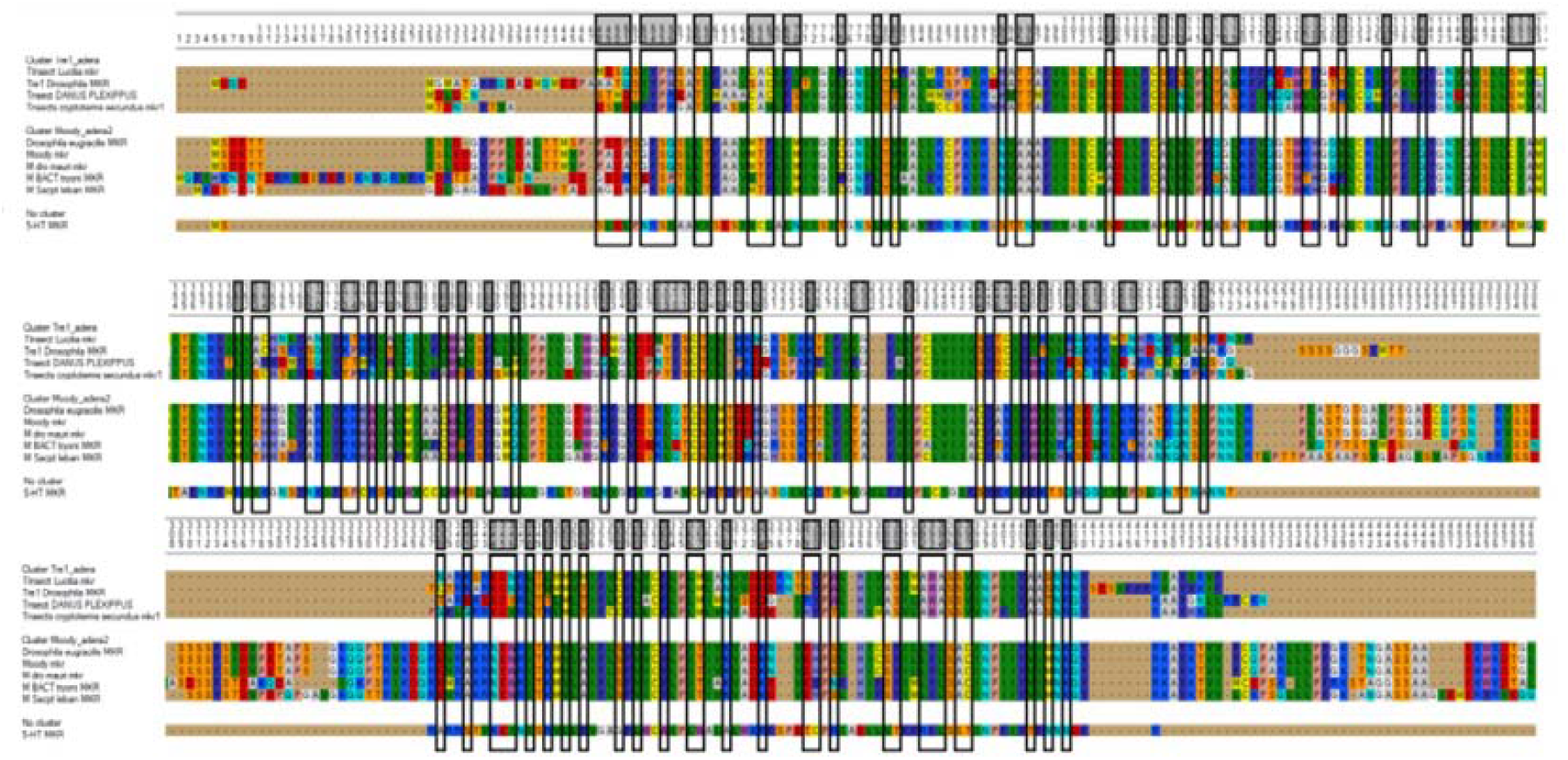
Functional dievrgence of the Tre1 family.

We only could locate Tre1 in any vertebrates, and CG319 was only confined to insects (figure 1). Interestingly Moody has been found in mice and humans (figure 1) along with cnidarian and epichem and hemichem. Thus Moody seems to have a continuous existence between invertebrates and vertebrates. Our positive selection analysis showed that Moody evolved under positive selection. This indicates that it could be various and additional function in higher vertebrates. However positive selection per branch was not significant. Taken together our results indicate that Tre1 super family diverged from an opsin like protein into Moody, later Tre1 appeared in Nephrozoa, while CG313 is only expressed in insects.

### Structural evolution and positive selection

The variation in the physiological functions of different groups of the Tre1 family could be influenced by possessing functional specific residues, functional divergent residues, being subjected to positive selection or a combination of these factors. It is interesting to note that Moody branch was subjected to statistically significant positive selection (Table 1). Type I functional divergence is the result of the change in evolutionary rate where a site is conserved for one group and is variable in another. We identified 1 critical functional divergence type I residues [add type 2]. Tre1 had an [K/, while Moody had a D/E/Q/R at the same location (which?). Intertsingly we found in the same location subsutuion of residues from [VTFS/RLQT]. This loctaion is inside the cell and not on the tramembrane or out. This may suggets that the main differnce ind unction between Tre1 homologs is related to thier downstream intecrtaion (evidecne).Motif search has shown that, Tre1 at position (37) Because crucial roles of Thr and Ser in membrane proteins have been proposed to be the formation of hydrogen bonds enhancing helix-helix interaction. In the same position in moody drosophila there exists (LL). Interestingly no Leucine appear at any identified functional diversion sites in Tre1. Conversely Leu also appears in other moody functional divergence sites such as 354. Moody is required for the maintenance of the Drosophila BBB. molecular replacement experiments revealed that the ruich lucine extracellular domain mediates binding to neurexins (Nrxs). In the fly BBB, glial cells establish intensive septate junctions that require the cell-adhesion molecule Neurexin IV. This observation indicate that moody could be enhancing BBB support through binding to Nrxs. Whereas, three functional divergence critical sites, appear in Tre1 and not in Moody, namely Met (position 127), Asp position (266) and Tyr (position 363). [add what is known about tre1 structure].

**Table 1.**
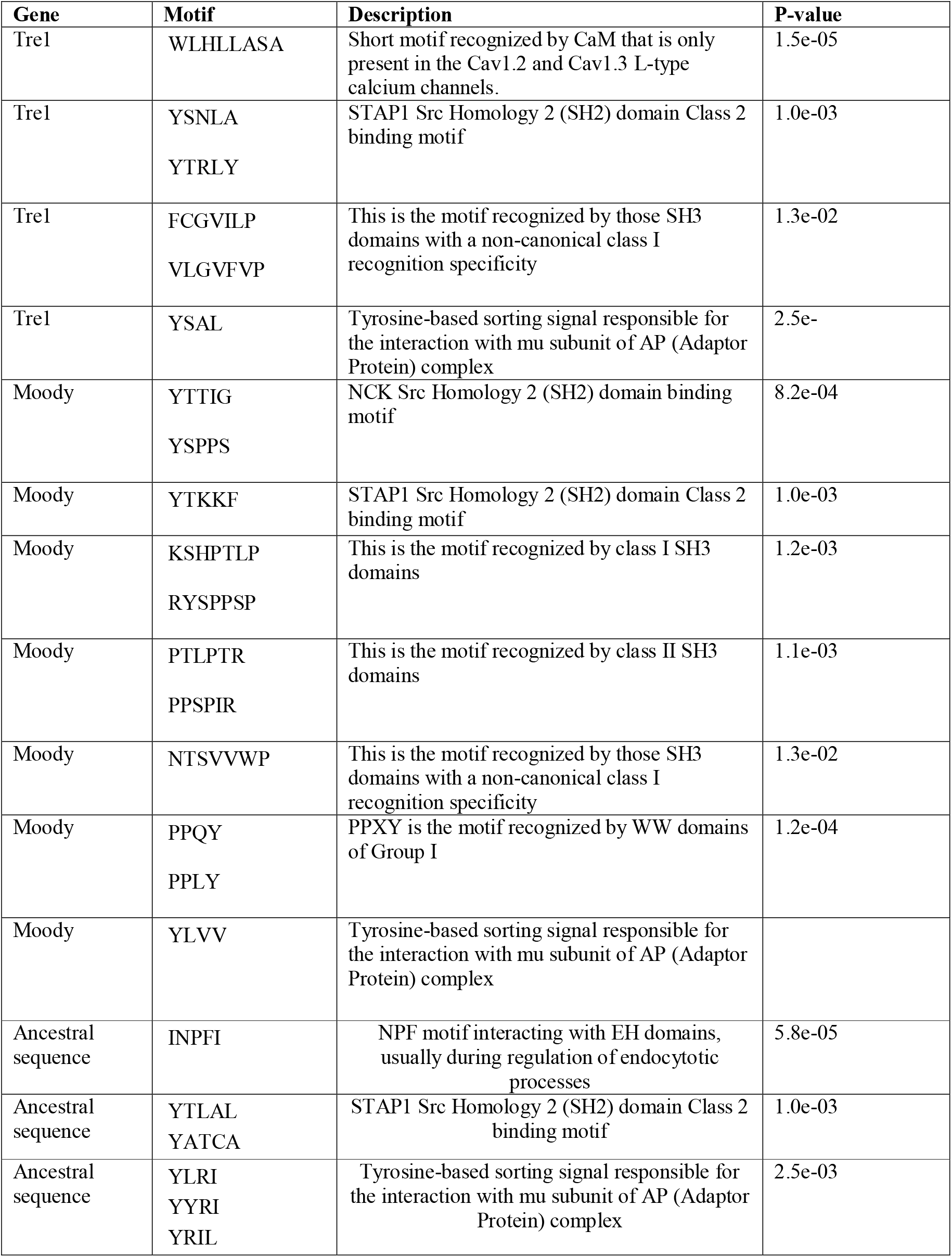
Motif shared by Tre1 family.

**Table 2.** Functional specficty motifs of the Tre1 family

|  |  |  |
| --- | --- | --- |
| outside | 1 | 37(35Thr) 36Leu |
| TMhelix | 38 | 60 |
| inside | 61 | 72 (72 thr) 72Ala |
| TMhelix | 73 | 95 |
| outside | 96 | 114 |
| TMhelix | 115 | 137( 126 ser,127 Met) 126Cys127Ile |
| inside | 138 | 156 |
| TMhelix | 157 | 176 |
| outside | 177 | 200(189Thr) 189Ser |
| TMhelix | 201 | 223(205 ile,206 Gly) 206Thr 207Ala |
| inside | 224 | 270(266Asp) 308Glu |
| TMhelix | 271 | 293 |
| outside | 294 | 302 |
| TMhelix | 303 | 325(314Ala) 354Leu |
| inside | 326 | 399(363Tyr) 584Pro |

**Table 3.** Functional divergence of Tre1 family in insects.

|  |  |  |  |
| --- | --- | --- | --- |
| outside | 1 | 14 |  |
| TMhelix | 15 | 37 |  |
| inside | 38 | 49 |  |
| TMhelix | 50 | 72 | [TE/LT][CAC/MTF] |
| outside | 73 | 91 |  |
| TMhelix | 92 | 114 | [TT/AA] |
| inside | 115 | 133 |  |
| TMhelix | 134 | 153 | [SMV/CIA] |
| outside | 154 | 177 |  |
| TMhelix | 178 | 200 |  |
| inside | 201 | 236 | [VTFS/RLQT] [K/DEQR] |
| TMhelix | 237 | 259 |  |
| outside | 260 | 268 |  |
| TMhelix | 269 | 291 |  |
| inside | 292 | 358 |  |

**Table 4.**
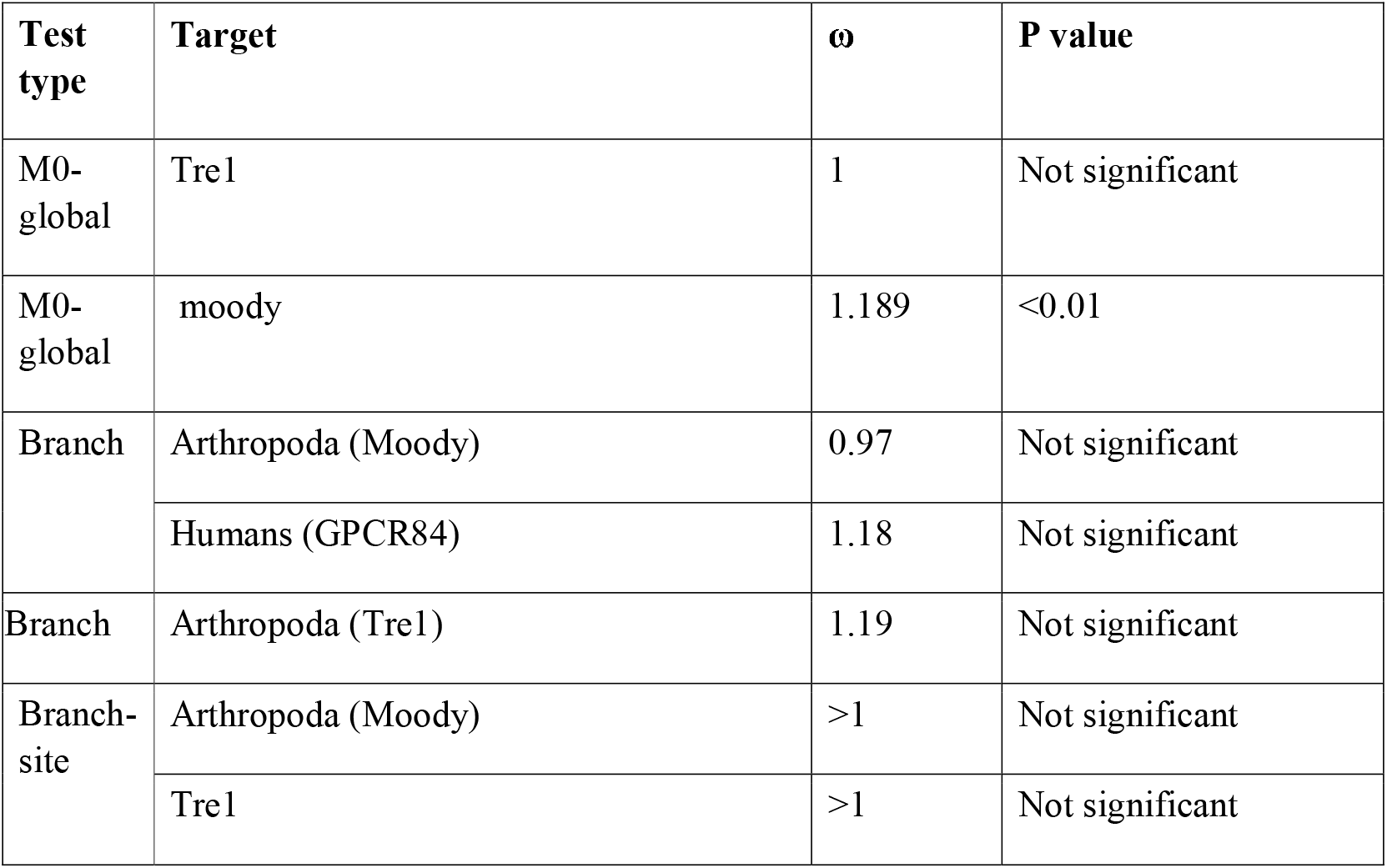
Likelihood values and parameter estimates for Tre1 genes under positive selection.

### Role of Tre1 and Moody in BBB

High similarity in evolutionary patterns, expression profiles, and structure between Tre1 groups hints at a possible role in brain morphological evolution in controlling BBB diapedesis. Drosophila has brain-like structure as well as immune cells-like known as hemocytes and BBB barrier like structure known as septal. The blood-brain barrier of Drosophila is established by surface glia, which ensheath the nerve cord and insulate it against the potassium-rich hemolymph by forming intercellular septate junctions. However, the diapedesis mechanism in drosophila is still unknown. In lower in vertebrates such as Trichoplax, no immune cells was found, and one rudimentary brain or neural cells were detected. Interestingly, CAV1 was found to be expressed in Trichoplax. CAV1 is likely to be used to construct caveolae for food transfer and its function has evolved to include cells later after the Cambrian explosion. This is supported by the observation that a rudimentary brain like structure has been reported in nematostella, with Two brain pathways Observed and immune-like cells known as ameboyctes. Only CAV1 has been detected and not PECAM1. In C. elegans, no immune cells were detected. However a glial BBB like structure was detected, together with CAV1. We as well as other have shown that CAV1 and PECAM1 play an indispensable role in BBB diapedesis ^21^. Interestingly, drosophila melanogaster does not express either CAV1 or PECAM1 which are vital for transcellular and paracellular diapedesis. Tre1 was reported to direct transepithelial migration of drosophila germ cells by regulating ecadherin^722^. It has also been shown to orient stem cell divisions in the Drosophila central nervous system^23^, control of germ cell polarity probably through controlling actin dynamics ^24 25^. Tre1 was also shown to be expressed in the drosphila BBB. However its function is not yet known. Additionally, moody controls blood-Brain Barrier Permeability in Drosophila^8^. It has been shown that Moody, the G protein subunits Gαi and Gαo, and the regulator of G protein signaling Loco are required in the surface glia to achieve effective insulation by regulating the cortical actin and thereby stabilizing the extended morphology of the surface glia. SH3 domains are small protein modules of about 50 to 60 residues that seem to be playing a part in the BBB integrity^26^. It is well known that SH3 through Rho pathway control actin movement in paracellular through PECAM1 mediated fashion^27 28^. Thus we could postulate that Tre1 and Moody could be playing a similar role in drosophila.

### Conclusion

The Tre1 family proposed time of divergence is during the time of radiation of Arthropoda, around 455[Mya. Our study identified several Tre1 residues as possible drug targets that could enhance controlling the BBB permeability. We were also able to reveal that NEGR1 might be interacting with the MAPK pathway as a form of signal transmitting receptor through its motif (KKVRVVVNF). In addition, we identified several amino acid residues that could be contributing to Tre1s’ axonal growth activity through controlling contact inhibition and metalloproteinase shedding.

